# A rebound from pre-dawn singing suppression drives intense morning singing in captive zebra finches

**DOI:** 10.1101/2025.09.29.679172

**Authors:** Ednei B. dos Santos, Chihiro Mori, Yunbok Kim, Joowon Kim, Ryosuke O. Tachibana, Ok-Yi Jeong, Juyeon Lee, Satoshi Kojima

**Affiliations:** Sensory and Motor Systems Research Group, Korea Brain Research Institute; Daegu, Republic of Korea; Department of Molecular Biology, Faculty of Pharmaceutical Sciences, Teikyo University; Tokyo, Japan; Department of Anatomy, Chonnam National University Medical School; Hwasun, Jeollanam-do, Republic of Korea; Human Informatics and Interaction Research Institute, National Institute of Advanced Industrial Science and Technology; Tsukuba, Japan; Department of Brain Sciences, Daegu Gyeongbuk Institute of Science and Technology; Daegu, Republic of Korea

## Abstract

Many birds sing vigorously at dawn, a phenomenon widely known as the “dawn chorus.” While its adaptive functions have been extensively studied, the proximate mechanisms underlying this widespread phenomenon have remained largely unexplored. Here, we demonstrate that intense morning singing resembling the dawn chorus is evoked as a rebound from singing suppression prior to dawn in captive zebra finches. Our findings indicate that when birds become hormonally aroused in darkness well before dawn, the suppression of spontaneous singing by darkness enhances their intrinsic motivation to sing. This elevated motivation subsequently triggers a rebound of intense singing immediately upon an increase in ambient light. These results reveal mechanisms by which intense morning singing arises from the interaction between the bird’s arousal state and ambient light, providing a plausible mechanistic explanation for the dawn chorus. Additionally, we found that rebound morning singing accelerates morning changes in song structure, supporting the existing hypothesis that the dawn chorus functions as a vocal warm-up to rapidly optimize song performance following nighttime silence. Together, our findings provide both mechanistic and functional insights into the dawn chorus and other diel vocal patterns in songbirds.

## Introduction

Animal vocal behavior is temporally structured rather than randomly distributed across the day. Such diel vocal patterns have been documented in diverse taxa, including primates^1,2^, frogs^3^, insects^4^, and birds^5^. Although these patterns are generally regulated by endogenous circadian mechanisms, they are also shaped by various extrinsic factors, such as light conditions and social contexts. It remains unclear, however, how endogenous circadian mechanisms interact with these extrinsic factors to orchestrate diel vocal patterns.

The “dawn chorus,” the intense singing of many bird species early in the morning, is perhaps one of the most conspicuous expressions of diel vocal patterns. It is typically defined as a period of elevated singing activity around dawn, beginning at twilight and peaking within a narrow temporal window^5,6^. In contrast to the extensive studies on the adaptive functions (i.e., ultimate causes) of the dawn chorus, the proximate mechanisms regulating when dawn singing begins and how strongly it is expressed remain largely unexplored. While the timing and intensity of the dawn chorus have been associated with various factors, including the sleep-regulating hormone melatonin, ambient light levels, social conditions, food availability, and reproductive state^5–7^, it remains largely unclear to what extent and how these factors interact to elicit intense dawn singing.

To elucidate the proximate mechanisms of the diel vocal patterns, including the dawn chorus, it is essential to investigate how individual external factors influence singing behavior by experimentally manipulating each factor independently. A recent field study utilized a solar eclipse to temporarily reduce ambient light levels during the daytime, and reported intense vocal activity resembling the dawn chorus following the temporary darkness in multiple avian species^8^. These findings suggest that temporary darkness alone is sufficient to trigger intense singing. Consistent with this, a previous study from our laboratory demonstrated similar, darkness-induced intense singing in captive zebra finches (*Taeniopygia guttata*)^9^, a widely used model organism in life sciences^10^. We found that in adult male zebra finches housed individually in sound-recording chambers, temporary lights-off during the daytime suppresses spontaneous non-courtship singing (referred to as “undirected singing”) and subsequently elicits a transient burst of intense singing when the lights are turned back on^9^. The latency and magnitude of this suppression-driven singing strongly depend on the duration of suppression, suggesting a mechanism in which the intrinsic motivation to sing accumulates while singing is suppressed, leading to a rebound of intense singing once the suppression is lifted. Given the striking parallel between eclipse-induced intense singing in wild birds and lights-off-induced intense singing in captive zebra finches, suppression-driven rebound singing may represent a common mechanism contributing to the diel vocal patterns across avian species.

Although daytime darkness induces rebound singing in both wild and captive birds, it remains unclear whether nighttime darkness similarly drives such a rebound in the morning. Since birds are typically asleep at night and sleeping birds do not exhibit singing behavior, singing suppression and resulting rebound singing are likely to depend on the birds’ arousal state as well as ambient light levels. Thus, the relative timing of bird’s awakening and morning light may be a critical factor determining the occurrence, onset, and intensity of rebound singing in the morning. Here, we test this prediction by shifting morning lights-on time and examining its effects on singing behavior in captive zebra finches. Our results demonstrate that intense morning singing can arise through a suppression-driven rebound mechanism, depending on the time lag between birds’ awakening and morning light, as predicted. We also provide evidence that intense morning singing occurs even under natural light conditions and may be mediated by similar rebound mechanisms. Additionally, we investigate the functional role of this intense morning singing by testing the hypothesis that intense morning singing serves as vocal warm-up to rapidly optimize song performance following nighttime silence^11–13^. We then discuss the implications of these findings in captive zebra finches for understanding the mechanisms and functions of the dawn chorus in wild birds.

## Results

### Captive zebra finches exhibit intense morning singing when morning lights-on time is delayed

Male zebra finches housed alone in sound recording chambers spontaneously produce hundreds of undirected songs per day under light conditions, but virtually no songs under complete darkness^14^. Our previous study demonstrated that temporary (5-10 h) suppression of undirected singing by turning off the lights during the daytime induces intense singing immediately after the lights are turned on and the suppression is lifted^9^. We first examined whether similar dark-induced intense singing would occur even in the morning following night-time darkness depending on the morning lights-on time (LT). We randomly shifted the morning LT by 3 hours earlier or later each day, relative to the regular LT, and examined their effects on undirected singing activity (Fig. 1A). Given that the magnitude of dark-induced singing during the daytime strongly depends on the duration of the dark period^9^, we predicted that delaying or advancing the morning LT (i.e., extending or shortening the night period) would enhance or attenuate morning singing activity, respectively. Although birds exhibited slight increases in singing rate relative to baseline even after the regular LT (0h LT in Fig. 1A-D & S1), these increases were markedly enhanced when the LT was delayed by 3 hours (+3 h LT in Fig. 1A–D & S1, Movie S1), resulting in intense morning singing. In contrast, almost no transient increases in singing rate were observed when the LT was advanced by 3 hours (−3 h LT in Fig. 1A–D, Movie S2). Moreover, birds began singing sooner after the delayed LT than after the advanced LT (Fig. 1E). These results demonstrate that intense morning singing resembling the dawn chorus can be induced in captive songbirds simply by delaying the morning LT.

**Fig. 1.**
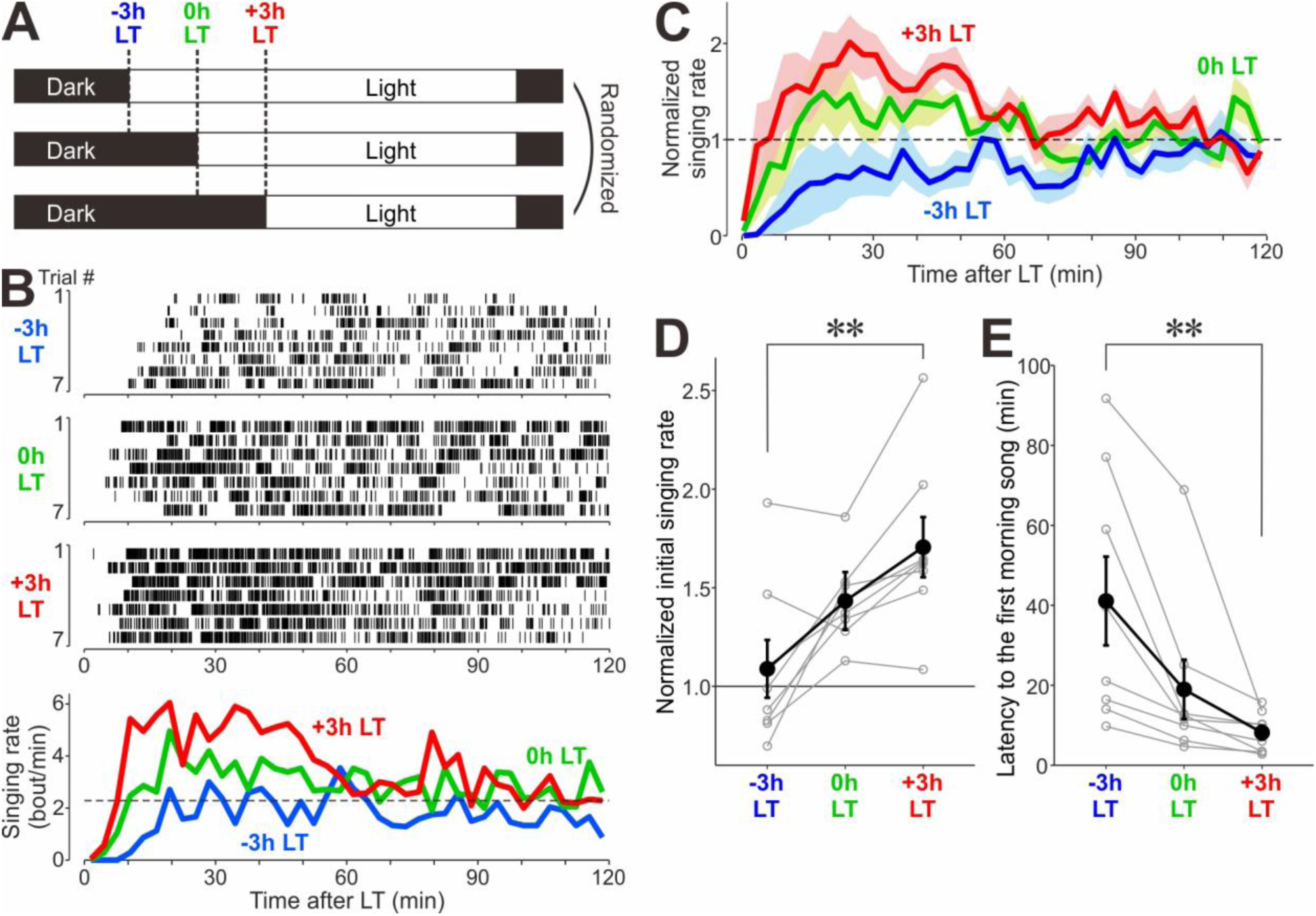
dawn chorus-like intense singing can be induced in captive zebra finches by delaying the LT in the morning. **(A)** Daily schedules of light and dark periods with variable LT (-3h LT, 0h LT, and +3h LT). The 3 LT conditions were randomized on a daily basis. **(B)** Raster plots of song bouts (*top*) produced after -3h LT, 0h LT, and +3h LT, and corresponding singing rate histograms (bin size, 3 min) (*bottom*) in a representative bird. **(C)** Time course of instantaneous singing rate in the three different LT conditions (normalized to the baseline rate, mean ± SEM, n = 8 birds). **(D)** Initial singing rate in -3h, 0h, and +3h LT conditions (normalized to baseline singing rate); gray circles with lines indicate the data of individual birds (*p* = 0.005 for 3 conditions, Friedman’s test; \*\**p* = 0.005 for -3h LT vs. +3h LT, Bonferroni corrected Durbin-Conover test). **(E)** Latency to the first morning song in 3 LT conditions (*p* = 0.0003 for 3 conditions; \*\**p* = 0.005 for -3h LT vs. +3h LT).

### Birds begin to sing in dimmer light conditions when LT is delayed

In addition to intense singing in the morning, the early onset of singing under dim light at dawn is another key feature of the dawn chorus in wild birds, and the onset of such dim light singing has been shown to be associated with the light input to the birds^5,6^. We found that singing under dim light conditions can also be induced by delaying the morning LT in captive zebra finches. We modified the morning lighting paradigm to mimic natural dawn conditions: Instead of the abrupt lighting used in the previous experiment, the light level was gradually increased over a 2-h period (see Methods), and this “gradual lighting period” was either delayed or advanced by 3 h relative to the regular LT (Fig. 2A-B). When the gradual lighting period was delayed by 3 h (+3h LT), birds began singing earlier within the gradual lighting period compared to when the gradual lighting period was advanced by 3 h (-3h LT) (Fig. 2B-D), indicating that singing initiation occurred under lower light levels when the morning LT was delayed. Taken together, these results suggest that delaying the LT can induce both elevated singing activity and singing under dim light conditions.

**Fig. 2.**
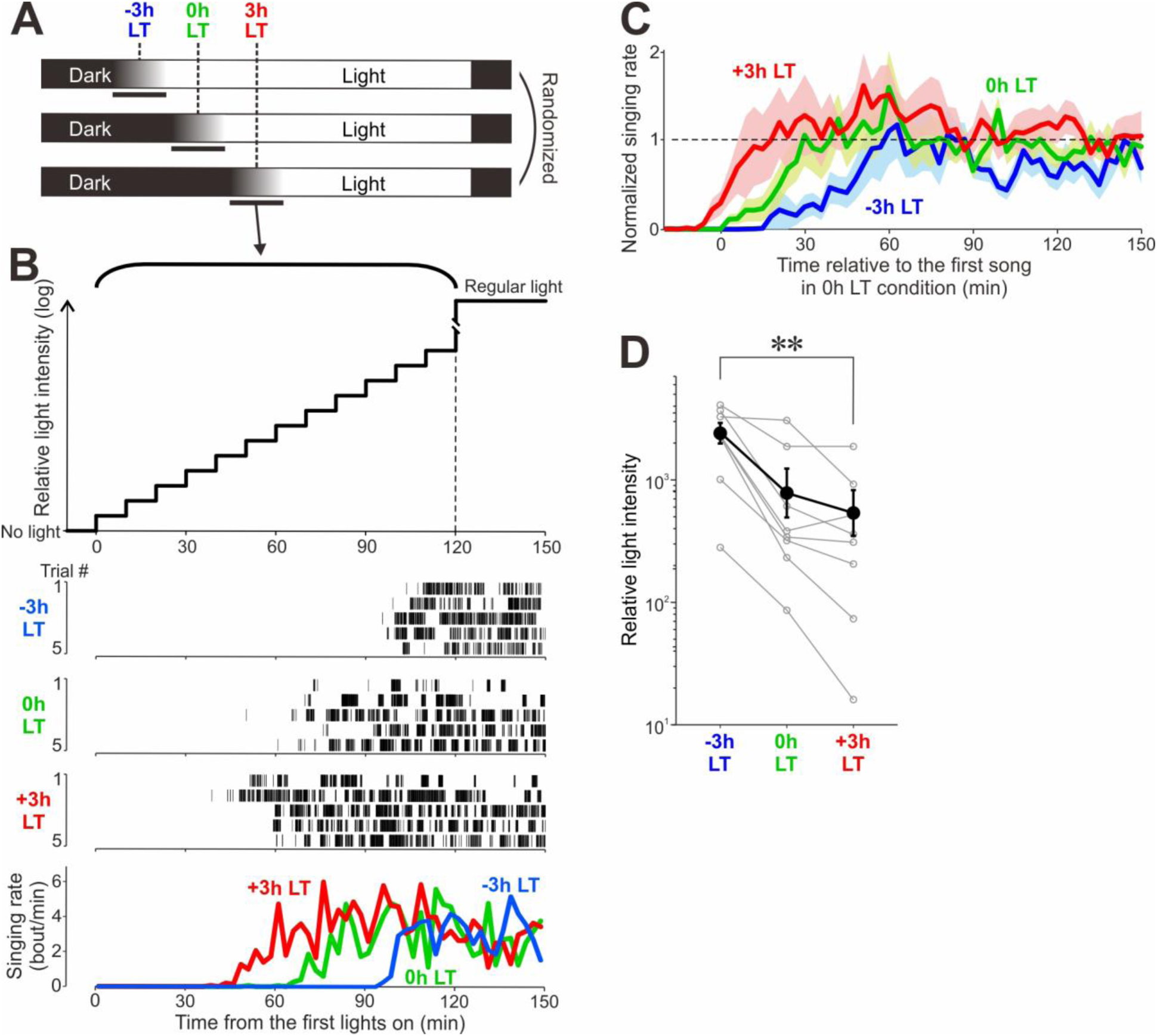
Birds begin to sing in dimmer light conditions when LT is delayed. **(A)** Daily schedules of light and dark periods with gradual lighting. The 3 LT conditions were randomized on a daily basis. **(B)** Time course of light level changes during a gradual lighting period (*top*), raster plots of song bouts during gradual lighting periods in -3h LT, 0h LT, and +3h LT conditions (*middle*), and corresponding singing rate histograms (*bottom*) in a representative bird. **(C)** Time course of instantaneous singing rate in the three different LT conditions (normalized to the baseline rate, mean ± SEM, n = 8 birds). The horizontal axis represents the time relative to the first song in the 0h LT condition. **(D)** Relative light intensity of first song in -3h, 0h, and +3h LT conditions (*p =* 0.0008 for 3 conditions; \*\**p =* 0.008 for -3h LT vs. +3h).

### Birds wake up well before the delayed LT and their motivation to sing increases while spontaneous singing is being suppressed by darkness

Why do birds exhibit intense morning singing with early onsets when the LT is delayed? Given that a similar pattern of intense singing occurs following temporary singing suppression by lights-off during the daytime^9^, we hypothesized that birds are mostly awake during the dark period before the delayed LT and that their motivation to sing increases while singing is being suppressed by darkness, subsequently producing intense singing as a ‘rebound’ from the suppression. To test this idea, we monitored the birds’ movements in darkness before the LT (with the abrupt lighting paradigm described in Fig. 1A) using infrared cameras mounted above individual cages (Fig. 3A-B). We found that birds moved actively during the 3-h period prior to the delayed LT (+3h LT in Fig. 3C-E, Movie S3), but exhibited little movement before the advanced LT (-3h LT in Fig. 3C-E, Movie S4). These results provide evidence that birds are mostly awake during the dark period before the delayed LT but not before the advanced LT.

**Fig. 3.**
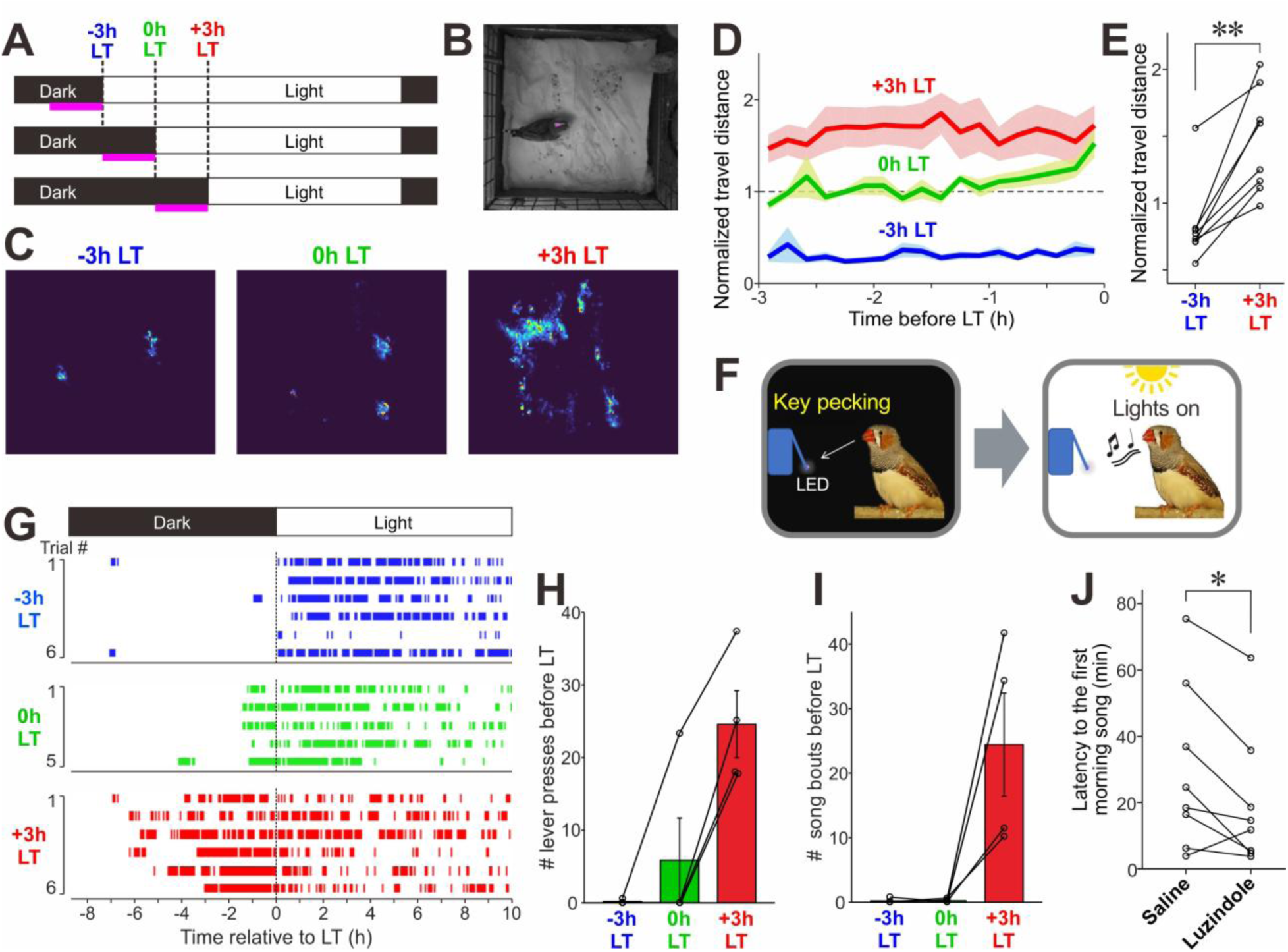
Birds are awake before the delayed LT and their intrinsic singing motivation increases while singing is being suppressed by darkness. **(A)** Daily schedule of light/dark periods and the 3-h periods of IR video recording (magenta lines). **(B)** Example IR video image of a bird in the dark. Magenta dot on the bird head indicates the point used to track bird’s movement. **(C)** Representative images indicating the positions of a bird during the 3-h dark periods immediately before the -3h, 0h, and +3h LTs (magenta lines in *A*). **(D)** Travel distances over the pre-LT 3-h periods (mean ± SEM across birds, normalized to the mean of 0h LT data for each bird). **(E)** Mean travel distances during the pre-LT 3-h periods, normalized to the 0h LT data, in individual birds (\*\**p* = 0.0078, Wilcoxon signed-rank test). **(F)** Schematics of operant conditioning paradigm to associate a lever press with a 10-sec period of lighting. **(G)** Representative plots of lever press activities (vertical bars) before and after the 3 different LTs. (**H & I)** The number of lever presses (H) and song bouts produced (I) before the 3 different LTs; lines represent the data of individual birds (averaged across trials) (*n* = 4 birds, *p* = 0.0324 and 0.038 for *H* and *I*, respectively, Friedman’s test). **(J)** The effect of the selective melatonin receptor antagonist luzindole on first song latency. Note that birds started singing significantly earlier when injected with luzindole compared to saline (\**p* = 0.027, paired *t-*test).

Although these results suggest higher arousal levels of the birds before the delayed LT, it does not necessarily indicate that they are highly motivated to sing at that time. To assess the singing motivation in the dark period before the LT more directly, we trained the birds to press a lever that triggered a 10-sec period of lighting (Fig. 3F, Movie S5). Since zebra finches sing almost exclusively under light conditions and not in darkness^14^, the frequency of lever presses is likely to reflect, at least to some extent, the levels of singing motivation as well as the arousal levels. When the LT was delayed or advanced by 3 h with the abrupt lighting paradigm, birds frequently pressed the lever and sang before the +3h LT, but only rarely before the -3h LT (Fig. 3G-I). These results support the idea that birds are highly motivated to sing in the dark before the delayed LT but not before the advanced LT. Given that suppression of undirected singing by darkness during the daytime increases intrinsic singing motivation and results in intense rebound singing immediately after the suppression is lifted^9^, these results provide evidence that intense morning singing is driven as a rebound from dark-induced singing suppression before the delayed LT.

We also investigated the hormonal mechanisms underlying early awakening before the delayed LT. In zebra finches, plasma concentrations of melatonin, a hormone that regulates the sleep-wake cycle in vertebrates^15^, are relatively high during the night and drop sharply to daytime levels approximately 0.5–2 hours before the regular LT^16^. This pre-dawn drop in melatonin levels roughly coincides with the early awakening of birds before the delayed LT. Based on these findings, we hypothesized that birds wake up well before the delayed LT due to hormonal regulation including the early melatonin drop but are suppressed from singing by darkness, leading to an increase in singing motivation and subsequent intense singing immediately after the LT. To test this hypothesis, we systemically administered the selective melatonin receptor antagonist luzindole 5 h before the regular LT and assessed whether this would induce intense morning singing. We found that birds began singing earlier after luzindole administration compared to saline administration (paired *t*-test: t(7) = 2.79, *p* = 0.0269; Fig. 3J), although the initial singing rates following the LT did not significantly differ between the two treatments (t(7) = 1.162, *p* = 0.284). These results suggest a key role of the early drop in plasma melatonin levels prior to the LT in inducing intense morning singing, thus supporting our hypothesis that the pre-dawn awakening that is responsible for intense morning singing is caused by melatonin-related hormonal regulation.

### Captive zebra finches housed under natural light conditions exhibit intense morning singing

Because our experiments so far were conducted in birds housed individually in sound recording chambers with artificial lights, it remains unclear whether similar intense singing is induced by natural morning light. To address this issue, we recorded the vocal activity of male and female zebra finches housed in an aviary exposed to natural light through large windows (over a 20-day period from mid-September to early December in 2022; the birds were housed with conspecifics with the same sex; see Methods for more detail). In these birds kept under natural light and social conditions (with almost constant temperature and humidity), the onset of their morning vocal activity appeared to vary depending on sunrise time and weather conditions (Fig. 4A), suggesting that light levels in the aviary critically influence morning vocal behavior. To determine whether these birds exhibited intense vocal activity in the morning, we accounted for variations in sunrise time and weather conditions by aligning the vocal activity data to a "dawn reference time (DRT)," defined as the time when light levels in the aviary surpassed a predetermined threshold (see Methods). This analysis revealed a marked increase in vocal activity around the DRT, followed by a gradual decline (Fig. 4B–C), resembling the typical dawn chorus observed in wild birds. Furthermore, detailed analysis of this morning vocal activity showed a relationship between dawn timing and the onset of morning vocalizations consistent with our findings in birds under artificial lighting (Fig. 1 & 2). Specifically, during the first 10-day recording period, in which both DRT and sunrise occurred relatively early, the average vocal activity tended to have a peak that was later than that during the subsequent 10-day period (Fig. 4B-C). Also, when the DRT occurred later due to seasonal changes and weather conditions, vocal activity around the DRT tended to be greater (Fig. 4D) and to start earlier (Fig. 4E). These findings in the birds under natural light and social conditions are consistent with the earlier onset and greater amplitude of morning singing after delayed LT observed in the birds under artificial light and socially isolated conditions (Fig. 1 & 2), suggesting similar mechanisms of morning vocal activity in the two conditions. These results provide evidence that natural morning light can induce intense vocal activity through the suppression-driven rebound mechanisms.

**Fig. 4.**
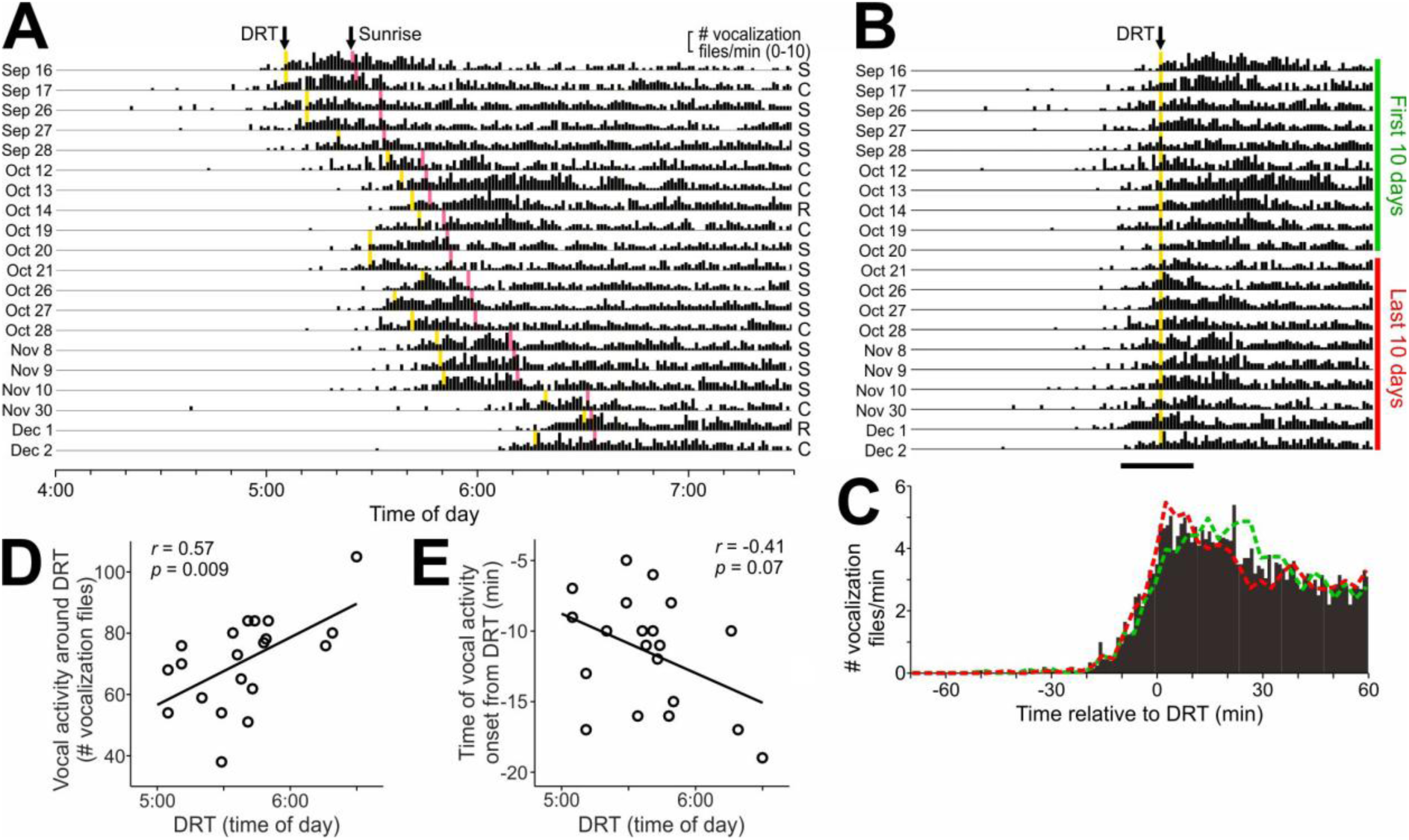
Captive zebra finches housed under social and natural light conditions exhibit dawn singing. **(A)** Vocal activity (number of vocalization files/min) recorded around dawn for 20 days from mid-September to early December. Yellow lines, dawn reference time (DRT); pink lines, sunrise time. Letters on the right side of the actogram indicate the weather on individual days: S, sunny; C, cloudy; R, rainy. **(B)** Vocal activity aligned to DRT; Green and red vertical lines indicate the first and last 10-day periods analyzed for the data in *C*. Horizontal bar at the bottom indicates the period in which vocal activity on individual days were measured for the plot *D*. **(C)** DRT-aligned vocal activity averaged over the all recording days (black histogram), the first 10-day period (green dashed line) and the last 10-day period (red dashed line). **(D)** Vocal activity during the 20-min period around DRT (horizontal bar in *B*) plotted against DRT; circles represent the data of individual days. Note that vocal activity around DRT was greater on days with later DRTs. **(E)** Time of vocal activity onset relative to DRT plotted against DRT. Note that vocal activity tended to start earlier relative to DRT on days with later DRTs.

### Intense morning singing accelerates morning changes in song structure

Our results so far have revealed the behavioral and hormonal mechanisms (i.e., the proximate cause) underlying intense morning singing in captive songbirds: In the birds that woke up well before LT, suppression of spontaneous singing by pre-dawn darkness increases singing motivation, leading to intense rebound singing shortly after the LT. We also investigated why this pre-dawn singing suppression increases singing motivation and induces rebound singing, thereby addressing the adaptive function (i.e., the ultimate cause) of intense morning singing. Recent studies on the dawn chorus in wild songbirds have demonstrated that song performance improves through dawn singing in a rate-dependent manner, leading to the "warm-up hypothesis" to explain the functional role of the dawn chorus^11–13^. This hypothesis posits that high-rate dawn singing serves as an intensive vocal warm-up that rapidly optimizes song performance to compensate for the lack of singing at night, providing an advantage in mate attraction and territory defense and ultimately increasing reproductive success. Since daily singing of captive adult finches has been repeatedly shown to serve as vocal exercise (or practice) to optimize and maintain song performance^17–24^, we wondered if the warm-up hypothesis might also apply to intense morning singing in captive zebra finches. To test this, we analyzed possible changes in song structure during morning singing in the songs recorded under the shifted LT conditions shown in Fig. 1, and examined whether intense morning singing accelerates such song changes.

We examined changes in morning song structure by comparing each song produced during the first hour after LT with all songs produced during the second hour in each LT condition (see Methods). Comparisons were made for each song syllable (n = 18 different syllables from 6 birds) by calculating mean acoustic distances in acoustic feature space (cosine distance, see Methods). In both the -3h LT and +3h LT conditions, many syllables showed gradual decreases in acoustic distance during the first hour (i.e., negative slopes of the acoustic distance trajectories; Fig. 5A-B), indicating substantial and monotonic changes in acoustic structure during morning singing. If these song changes were caused by the act of singing rather than simply by the passage of time, the speed of the song changes should depend on the singing rate during that period. Consistent with this prediction, we found that the slopes of song changes over the first 30 min were significantly steeper in the +3h LT condition, where intense morning singing occurred, than in the -3h LT condition (Fig. 5A-B and Table S1). Moreover, the slopes were significantly correlated with normalized singing rate during the same period (the number of syllable renditions in the first 30-min period normalized to that in the first 2-h period) (Fig. 5C and S2). These results suggest that morning song structure changed as a function of recent singing activity and that such changes were accelerated by intense morning singing. This conclusion was further supported by the findings that, when acoustic distances were plotted as a function of syllable order instead of time, there was no significant difference in the slopes between the -3h and +3h LT conditions (Fig. 5D-E and Table S2). In summary, these findings are consistent with the warm-up hypothesis for the dawn chorus that intense morning singing functions as a vocal warm-up that rapidly optimizes song performance following overnight silence.

**Fig. 5.**
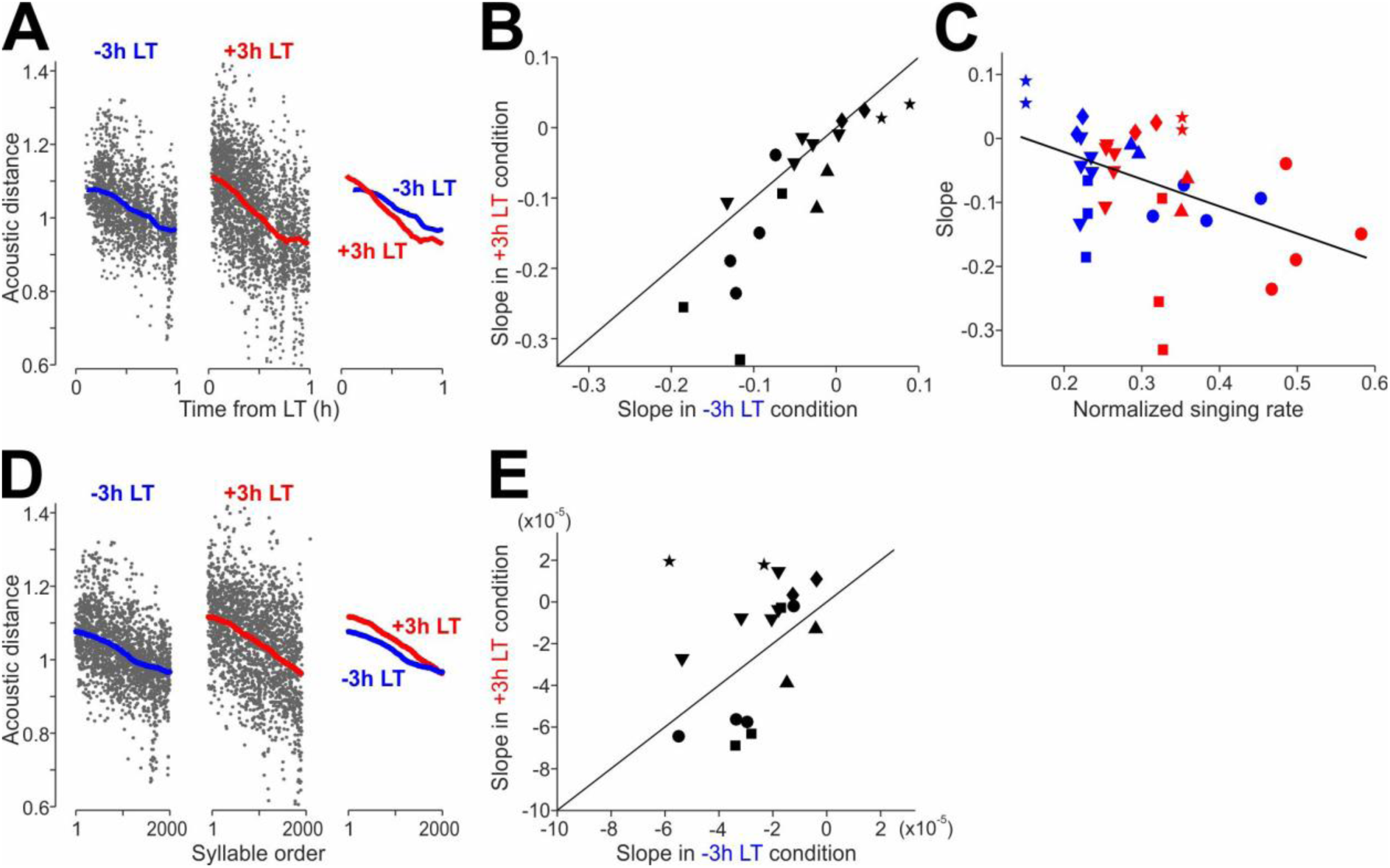
dawn singing accelerates morning changes in song syllable structure. **(A)** Changes in acoustic distances of syllable structure in the renditions during the first hour after LT relative to the renditions during the second hour in -3h and +3h LT conditions for an example syllable. Gray dots and thick lines indicate the data of individual syllable renditions and their local mean trajectories, respectively; the local mean trajectories in the 2 conditions are overlaid in the rightmost plot. **(B)** Comparisons of the slopes of acoustic distance changes over the first 30 min between -3h and +3h LT conditions (n = 18 syllable types from 6 birds, t = 2.74, *p* = 0.0140, a linear mixed-effects model including random intercepts for bird identity and syllable; see Table S1). Each data point indicates a data from individual syllable type; different symbols represent different birds; Diagonal line indicates unity. **(C)** The same slope data in the 2 LT conditions plotted against the singing rate during the first 30-min period (the singing rate was normalized to that in the first 2-h period). Blue and red symbols indicate data in -3h and +3h LT conditions, respectively; symbols are as in *B* (n = 36, pairwise linear correlation, ρ = -0.4445, *p* = 0.0066). **(D)** Acoustic distances of the same syllable as shown in A, plotted as a function of syllable order instead of time. Conversions are identical to those in A. **(E)** Comparisons of the slopes over the first 1000 syllable renditions between -3h and +3h conditions (t = 0.35, *p* = 0.3509, linear mixed-effects model with random intercepts for bird identity and syllable; see Table S2).

## Discussion

In the present study, we demonstrated that intense morning singing in captive songbirds is induced as a rebound from singing suppression caused by pre-dawn darkness. Under artificial light and socially isolated conditions, birds woke up in the dark well before the delayed dawn, likely mediated by hormonal mechanisms involving melatonin, and their intrinsic motivation to sing increased while spontaneous singing was suppressed by darkness. When the suppression was lifted by morning light, the elevated singing motivation triggered intense morning singing as a rebound from the suppression. Similar rebound-like vocal activity was induced by natural morning light in birds housed under social conditions. These results revealed mechanisms by which intense morning singing arises from the interaction between the arousal state of the birds and ambient light, and highlight rebound singing as playing a key role in regulating diel patterns of vocal activity in songbirds.

Although the intense morning singing observed in captive zebra finches resembles the dawn chorus generally observed in many avian species in the wild, further studies are needed to determine whether the rebound singing mechanism is applicable to the dawn chorus. Importantly, to our knowledge, no previous studies have reported a dawn chorus in wild zebra finches, and even the overall singing rate in a solo context in the wild (i.e., undirected singing) appears to be much lower than that in a laboratory setting^25^. The causes of these differences in singing behavior between the two conditions are unclear, but they may be due, at least in part, to the high predation risk in the wild, which would reduce solo singing, including rebound-driven intense singing at dawn. In addition, singing rates of male zebra finches in the wild vary depending on social context, in particular whether they are with their female partners, as well as on their breeding stage and season^25^. Given that our experiments were conducted on birds that were housed either in socially isolated or together with the same-sex conspecifics and thus were not in breeding conditions, it is also possible that overall singing activity, including dawn chorus-like intense singing, is largely reduced in the wild due to such differences in social context and breeding conditions. Other social and environmental factors, such as food availability, meteorological conditions, and the degree of freedom of movement, may also significantly affect their singing activity and attenuate intense morning singing.

Moreover, it is possible that, although zebra finches have the capability to produce suppression-driven rebound singing following temporary darkness both in the morning and during the daytime^9^, this trait may be unique to zebra finches and not found in other avian species. However, a similar transient increase in vocal activity has been reported following temporary darkness caused by a total solar eclipse in multiple avian species^8^, showing a striking parallel with rebound singing following temporary darkness in captive zebra finches^9^. Thus, it is reasonable to assume that rebound singing is a common trait of many avian species, including the zebra finch. Importantly, species that exhibit a natural dawn chorus are more likely to show the eclipse-induced intense vocal activity^8^, suggesting a potential link between dark-induced rebound singing and the dawn chorus. Given these findings and implications, we propose a novel hypothesis for the proximate mechanisms of the dawn chorus: a rebound from dark-induced singing suppression is a mechanism that can drive intense dawn singing in many bird species, but its expression critically depends on various social and environmental factors.

A key aspect of the rebound singing mechanism identified in the present study is the time lag between when birds wake up and when light intensity rises. During this time lag, birds are awake but suppressed from singing by darkness, and their intrinsic singing motivation appears to increase depending on the duration of the time lag, as in birds that are suppressed from singing by lights-off during the daytime^9^. This suppression-dependent increase in singing motivation is reminiscent of, and consistent with, the so-called “Lorenz’s psychohydraulic model,” in which a reservoir of motivational impulse builds up and is ultimately released by appropriate environmental stimuli^26^. Although mechanisms underlying this motivational increase remain largely unclear, a brain region and neuromodulators related to singing motivation have recently been reported^9,27,28^. On the other hand, the mechanisms by which birds awaken before dawn likely involve hormonal regulation including a drop in plasma melatonin concentrations from early night levels^16^, which is driven by a circadian clock even in the absence of light inputs^29,30^. A similar trend of decreased melatonin levels during the late-night phase has been reported in songbirds other than zebra finches, although the timing of these changes remains unclear^31–34^. Importantly, in wild songbirds, nighttime melatonin concentrations are positively associated with the onset time of early morning behaviors including dawn singing^33,35^, suggesting a possible link between the circadian secretion of melatonin and the dawn chorus^5,6^. On the other hand, the timing of the dawn chorus has also been suggested to be associated with light inputs to birds, such as the intensity of morning light and the size of their eyes^36–39^. Our findings of dark-induced rebound singing in the morning provide a plausible explanation to reconcile these seemingly contradictory findings, and propose potential mechanisms by which the circadian regulation of melatonin and light inputs interact to determine both the timing and magnitude of the dawn chorus. Future studies should investigate whether birds are awake long before dawn and if these arousal states are correlated with the timing and intensity of the dawn chorus.

In addition to addressing the proximate mechanisms of intense morning singing, our results provide insight into its potential functional significance. The acceleration of morning song changes by intense singing is consistent with the warm-up hypothesis previously proposed for the dawn chorus in wild birds^11–13^. This hypothesis posits that intense dawn singing serves as intensive vocal warm-up that rapidly optimizes song performance to compensate for the overnight lack of singing, ultimately enhancing reproductive success. This hypothesized function appears to be well matched with our finding that, when birds are awake well before dawn but suppressed from singing by darkness, their intrinsic singing motivation increases, leading to intense morning singing as a rebound from the suppression. Importantly, however, our data do not directly address whether such structural changes are functionally adaptive or preferred by females. Since our analysis focused on changes in song structure rather than performance evaluation or reproductive outcomes, further research is necessary to determine whether these changes constitute improvements with functional consequences in zebra finches. Nevertheless, several previous studies support the notion of song improvement. For example, in juvenile zebra finches undergoing song development, song structure deteriorates overnight and then dramatically improves (i.e., approaches their final adult song) through morning singing^40^. In addition, even in adult zebra finches, which normally produce songs with stable overall structure, song performance defined as the degree of female’s preference to that song significantly decreases by prolonged suppression of daily singing and gradually recovers after singing is resumed^17,18^. Thus, it is reasonable to suggest that our results of morning singing-dependent song changes reflect improvements in song structure.

Although it remains to be determined whether the rebound-based mechanism and the hypothesized warm-up function of intense morning singing apply to the dawn chorus observed in wild birds, these ideas do not necessarily contradict existing hypotheses about the dawn chorus. In fact, the rebound mechanism and the warm-up hypothesis can, at least in theory, explain many dawn chorus-related findings that support existing hypotheses. For example, previous studies have reported that the extent of the dawn chorus (e.g., singing duration and the timing of the first song) is associated with specific reproductive stages, such as the female fertility period, and is also positively correlated with reproductive success (see^5,6^ for review). Based on these findings, it has been proposed that the dawn chorus represents an honest signal of male quality^41–43^ and directly regulates female reproductive behavior around dawn^44,45^. The warm-up hypothesis states that the dawn chorus functions to rapidly optimize the quality of song performance, which is critical for mate attraction when produced in a courtship context. Thus, it is possible that birds exhibit a more pronounced dawn chorus during the female fertility period in order to optimize song performance as early as possible in the morning and to maximize mating success. The rapid song optimization by the dawn chorus could also be advantageous for territory defense in many territorial species. Although they need to defend their territories throughout the day, they may rapidly optimize their song performance through the dawn chorus early in the morning to prepare for the territory intrusion by other birds. This view is consistent with a previous study showing that the extent of the dawn chorus is affected by territorial intruders^46^ and also explains the observation that many songbird species exhibit the dawn chorus even after the female fertility period. Additionally, because birds exhibiting a more intense dawn chorus are likely to have higher motivation, not only for singing, but also for other reproductive behaviors, such elevated motivation for non-singing behaviors could also contribute to higher reproductive success. Moreover, our rebound singing hypothesis also aligns with the unpredictable overnight-condition hypothesis, which proposes that birds sing intensely at dawn to expend surplus energy stored in preparation for unpredictable overnight conditions^47^. This hypothesis is supported by findings that greater food availability leads to a more pronounced dawn chorus^48–50^. Since food availability positively affects birds’ nutritional state and overall singing motivation, as shown in captive zebra finches^27^, sufficient food availability could amplify the dawn chorus by elevating overall singing motivation and enhancing suppression-driven rebound singing.

In conclusion, the present study highlights suppression-driven rebound singing as playing a key role in shaping diel patterns of vocal activity in captive songbirds. Because rebound singing requires a condition in which birds are awake yet their singing is suppressed by darkness, its occurrence depends critically on the interaction between circadian-regulated arousal and ambient light. These findings lead to a novel hypothesis that can explain the proximate mechanisms of the dawn chorus, one of the most prominent diel patterns of avian vocal activity. As this mechanistic hypothesis aligns well with existing hypothesis on its adaptive functions, the present study opens up a new avenue in dawn chorus research, including experimental tests for the rebound mechanism in wild birds.

## Materials and Methods

### Subjects

The subjects used for song recording under socially isolated and artificial light conditions were adult male zebra finches (*Taeniopygia guttata*, 87–260 dph) bred in the songbird facility at the Korea Brain Research Institute (KBRI). They were raised with their parents and siblings until ∼60 dph and then housed with their siblings and/or other males conspecifics until the experiments started. Their care and treatment were reviewed and approved by the Institutional Animal Care and Use Committee (IACUC) at the KBRI. The subjects used for recording vocal activity under social and natural light conditions were adult male and female zebra finches bred in the songbird facility at the Teikyo University. Their care and treatment were reviewed and approved by the Animal Care and Use Committee there. All experimental procedures were carried out in compliance with the ARRIVE guidelines^51^.

### Song recording

For the experiments with song recording under socially isolated and artificial light conditions, each bird was housed in a sound-attenuating and light-proof chamber (MC-050, Muromachi Kikai, or custom-made chambers) throughout the experiment. Songs were recorded using a microphone (PRO35, Audio-Technica) positioned above the cage and a custom-written song recording program (tRec, R. O. Tachibana). Output from the microphone was amplified by a mixer (402-VLZ4, Mackie) and digitized via an audio interface (Octa-Capture UA-1010, Roland) at 44.1-kHz (16-bit). Recorded signals were down-sampled to a sampling rate of 32-kHz. Recording was triggered if the program detected 3 or 4 consecutive sound notes, each of which was defined based on sound amplitude, duration, and intervening gap duration; those parameters for triggering sound recording were optimized for each bird so as to record all songs produced. Each recording ended if a silent period lasted longer than 0.5 sec (i.e. each song file contains a single “song bout” that is separated from other bouts by >0.5-sec silent periods). Songs were recorded throughout the day, and all song recordings were of undirected song (i.e. no female was present). Birds with sufficient singing rates (>300 song bouts per day) were used for the experiments. The procedures to record vocal activity of the birds housed under social and natural light conditions and to analyze the recorded vocal activity are described below.

### Manipulation of light-dark schedule and illumination level

Birds for the experiments under socially isolated and artificial light conditions (Figs 1-3) were bred and kept on a constant 14:10hr light:dark (LD) cycle (with abrupt light-dark transitions) in the bird colony and after being transferred to the sound-attenuating chambers for > 5 days until the experiments began. During the experiments, we used two different lighting protocols. For the “abrupt lighting” experiment, the time of abrupt dark-light transition (the lights-on time, LT) was shifted daily ±3-h from the original 14:10hr LD cycle; birds underwent the -3hr, 0hr, and +3h LT conditions in a randomized order (neither the -3h LT nor the +3h LT condition was administered on consecutive days to prevent the birds from adapting to the LT condition). Each condition was administered multiple times (ranging from 4 to 8 trials). For the “gradual lighting” experiment, the illumination level was exponentially increased with 12 steps over 2 hr (10-min period for each step) using LED lights with an Arduino-based control unit; the illumination level of the 12^th^ step was approx. 350 lux, and it was followed by a constant illumination level equivalent to the one used for regular light periods (approx. 500 lux). This 2-hr gradual lighting period was shifted daily ±3-h in a randomized order just as for the abrupt lighting experiment. Illuminance was measured at the center of the experimental cages using a light meter (Tenmars, TM-205).

### Song analysis

To analyze singing activity recorded in sound-attenuating chambers, we screened all sound files recorded during the time periods of interest to exclude non-song files using a semi-automated method previously described^9^. Briefly, we sorted song files (sound files that include at least one full motif of song) and non-song files by focusing on temporal trajectories of sound amplitude (amplitude envelopes) and those of Weiner entropy, both of which are highly stereotyped across motif renditions and clearly distinct from other sounds including non-song vocalizations. We calculated the maximum correlation coefficients (mCCs) of amplitude envelopes and that of entropy trajectories between a canonical song motif and all sound files recorded, and excluded files that had mCCs for both amplitude envelopes and entropy trajectories below certain thresholds as non-song files. We then visually inspected spectrograms of those excluded files to ensure that no song files were excluded from the song dataset to be analyzed.

To quantify the magnitude of dawn chorus-like singing activity, we used two measures, the initial singing rate and first song latency^16^. The initial singing rate was measured as the mean singing rate over a 1-hr period starting at the onset of the first song produced in the morning for each trial and averaged across trials with the same LT. The initial singing rate was then normalized to the baseline singing rate, which was cross-trial and cross-LT average of the mean singing rate over the 30-min period starting at 90 min after the morning LT. We also plotted time courses of singing activity by measuring instantaneous singing rates using 3-min bins and averaged across trials with the same LT. The average instantaneous singing rates were then normalized by the baseline singing rates. The first song latency was measured in the abrupt lighting experiments as the time interval from the LT to the onset of the first song recorded in the morning. The latencies were then averaged across trials with the same LT.

To investigate morning changes in song structure, songs recorded in the -3h LT and +3h LT conditions were segmented into syllables using a custom program on Matlab based on the following parameters: amplitude, minimum and maximum syllable duration, and minimum and maximum frequency. These parameters were manually adjusted for each bird. From individual syllables, 8 time-varying acoustic features (centroid, spread, skewness, kurtosis, flatness, slope, pitch, goodness of pitch) and 2 other features (amplitude and duration) were extracted and projected onto a two-dimensional space by dimensionality reduction algorithm t-SNE (t-distributed Stochastic Neighbor Embedding). Syllables were then manually labelled based on clusters formed on the two-dimensional space. All labeled syllables were manually checked and incorrect labels were corrected. For the syllables that had stereotyped temporal structure across renditions, morning changes in acoustic structure were examined in each LT condition by comparing each syllable rendition produced during the first hour after LT with all syllable renditions produced during the second hour (the reference period) and by averaging their acoustic distances. The acoustic distance was quantified as cosine distance computed using the within-syllable mean, standard deviation, and mean absolute derivative of the 8 acoustic features and amplitude described above. The acoustic distances of individual syllable renditions were plotted as a function of time after LT and the local means over a sliding window of ±10 min durations were computed with 1-min increments. Slopes of the local mean changes over the first 30-min period starting at the first rendition were compared between the -3h and +3h LT conditions. The same acoustic distance data were also plotted as a function of syllable order instead of time, and slopes of the local mean changes were compared between the 2 conditions. Additionally, the slopes of acoustic distances were plotted against the number of syllable renditions recorded during the first 30-min period normalized to the number of renditions recorded during the first 2-h period to examine correlations between them.

### Recording and analysis of bird movements

In the birds that underwent the ±3h LT conditions with the abrupt lighting paradigm, their movements in the dark before the LT were monitored using infrared cameras (SQ12, Mrs Win, and UI-3040CP Rev. 2, iDS imaging) positioned on the top of the individual cages. Infrared LED lights were also positioned ∼10 cm above the cages and they were on during the night dark period. Each recording session spanned 3 hours, finishing at the LT (-3h, 0h, or +3h LT). To increase the precision of movement tracking, a small fluorescent sticker was affixed on the top of each bird’s head. Movements of the birds’ heads were tracked using the machine learning software DeepLabCut^52^, and their travel distances were calculated over 10-min time bins. Mean travel distances over the 3-h period were calculated for each LT condition and compared across different LT conditions.

### Operant lever-pressing task for lighting

To examine the level of motivation in the dark to seek light conditions that allow birds to sing, we trained birds to press a lever that is associated with the activation of ambient light. Birds were kept individually in an experimental cage that was placed in a sound-attenuating and light-proof chamber throughout the experiments. On the ceiling of the chamber, white LED lights were installed in addition to the built-in lights of the chamber. In the cage, a lever switch (Omron Electronics, SS-01GL111-E) was placed on the rear wall near a perch, and a small red LED was also placed on the rear wall above the lever. Throughout the experiment, including day and night, pressing the lever immediately activated the white LED lights for 10 seconds, and the timing of lever pressing by the bird was recorded; the small red LED was always lit so that the lever location could be identified in the dark. Undirected songs were also recorded throughout the experiment using a microphone positioned above the cage. Birds were initially housed in the experimental cage with the lever operational on a regular 14/10 light-dark (LD) cycle for more than 7 days for the purpose of habituating the bird to the experimental apparatus and having them to learn the association between lever pressing and activation of white LED lights. Subsequently, we introduced the behavioral paradigm of shifting the morning LT described above (the “abrupt lighting experiment”): birds underwent -3hr, 0hr, and +3h LT in a randomized order. Each condition was administered multiple times (ranging from 4 to 7 trials). The frequencies of lever pressing and of song bouts over the 3-h period before the LT were averaged across trials and compared across different LT conditions.

### Luzindole injections

A potential role of melatonin in morning singing was examined using luzindole (Tocris, 0877), a competitive melatonin MT1/MT2 receptor antagonist^53^. Luzindole was stored as stock solution in DMSO (40 mg/ml) at −20 °C and diluted in PBS before use (2 mg/ml). Birds were kept on a regular 14/10 LD cycle with abrupt light-dark transitions, and injected subcutaneously with luzindole (5.0 mg/kg) or PBS 5 h before the LT. Each bird received 4 injections for both luzindole and PBS in a randomized order (a single injection per day). Undirected songs were recorded throughout the experiment, and morning singing was analyzed by focusing on the first song latency and initial singing rate. Results were averaged across trials for each bird.

### Recording and analysis of vocal activity of the birds under social and natural light conditions

To examine morning vocal activity under social and natural light conditions, audio and video recordings were made in the zebra finch aviary at Teikyo University, which has large glass windows that let in natural light from outdoor, for 20 days intermittently from September 16th to December 2nd, 2022. In the aviary, ∼90 zebra finches were housed in 18 cages (8 cages for breeding pairs and 10 cages for adult birds with the same sex) that were placed in a metal rack (4 cages x 3 tiers) near the windows. Audio recordings were made using a microphone (PRO35, Audio-Technica) placed in the middle of the cage rack and connected to an audio interface (Octo-Capture UA-1010, Roland) for digitization at 44.1 kHz (16 bit). To capture vocal sounds, we used a custom-made recording program, which started sound recording when it detected 5 successive sound notes, each of which reached predefined thresholds of amplitude and duration, and ended when it detected a silent period lasted >0.8 sec. This recording system captured mostly vocal sounds (i.e., songs and calls), but non-vocal sounds such as cage noises and wing-flapping sounds were occasionally recorded. To measure vocal activity around dawn, we visually sorted the files that contain vocalizations using Avisoft Saslab (Avisoft Bioacoustics), and calculated the number of the files recorded during 1-min time bins over the time period from 4:00 to 7:30 am. The daily vocal activity data were then aligned relative to the “dawn reference time (DRT)”, which was defined as follows: During the periods of audio recordings, we simultaneously videotaped all cages and windows in the aviary using a wide-angle web camera (BSW200MGK, Buffalo) placed 80 cm away from the cage rack (frame rate, 16 Hz). On the recorded videos, the minute-by-minute changes in luminance for four window locations (10 x 10 pixels) were measured using Fiji (ImageJ, version 1.54f^54^). For each day, the time point when the luminance reached 200 was defined as the DRT, and the vocal activity data were aligned to the DRT. The aligned data were then averaged across days to obtain the average vocal activity patterns. We obtained sunrise times and whether conditions for individual experimental days at the website of the National Astronomical Observatory of Japan (https://eco.mtk.nao.ac.jp/cgi-bin/koyomi/koyomix_en.cgi) and that of the Japan Weather Association (https://tenki.jp/past/2022/09/weather/).

### Statistics

Statistical tests were performed using MATLAB (MathWorks) or R (R Core Team and R Foundation for Statistical Computing). To the differences in singing and lever-press activity across different LT conditions, we used a Friedman’s test followed by Bonferroni corrected Durbin-Conover test. To examine the effects of different LT conditions on birds’ travel distances and the effects of luzindole administration on morning singing activity, we used a Wilcoxon signed-rank test and a paired t-test, respectively. To assess the effects of different LT conditions on the slopes of morning song changes, we fitted a linear mixed-effects model predicting each syllable’s slope from LT condition (–3h LT vs +3h LT), while accounting for two levels of nesting: each bird has its own baseline (random intercept) and, within each bird, each syllable has its own baseline (another random intercept). A p-value of <0.05 was used as the criterion for statistical significance.

## Acknowledgments

We thank D. Rendall at University of New Brunswick, D. M. Logue at University of Lethbridge, J. Sakata at Mcgill University, and N. Rattenborg at Max-Plank Institute for discussion and comments on this manuscript.

## Supplementary Materials

Materials and Methods

Figs. S1 to S2

Tables S1 to S2

References (50–53)

Movies S1 to S5

