## Supplementary material for "A rebound from pre-dawn singing suppression drives intense morning singing in captive zebra finches": Figs. S1 to S2 and Tables S1 to S2

**The PDF file includes:**
Figs. S1 to S2
Tables S1 to S2
References (*50–53*)

**Other Supplementary Materials for this manuscript include the following:**
Movies S1 to S5

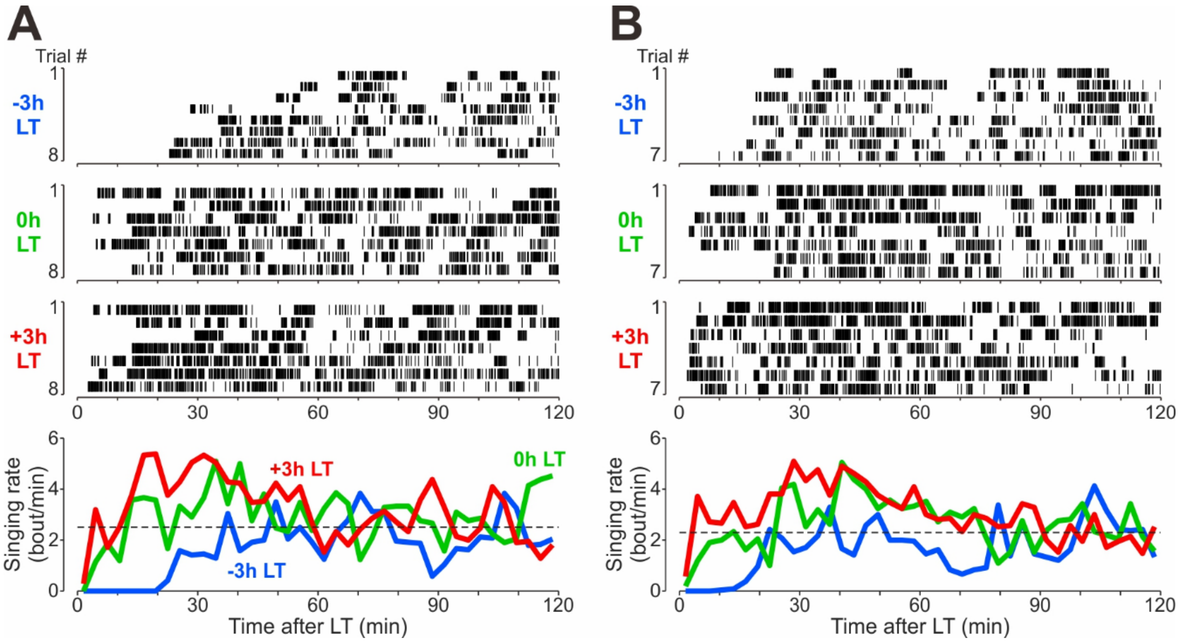


Fig. S1. LT-dependent DC-like singing in two addition birds. Conventions are as in Fig. 1B.

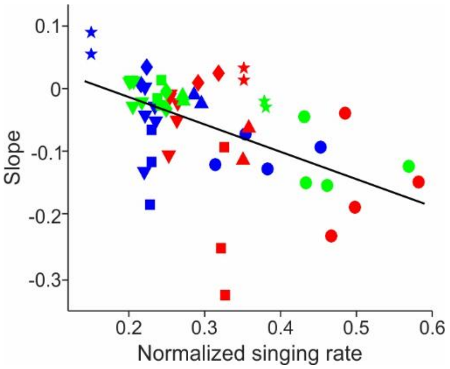


Fig. S2. The slope data in -3h (blue) and +3h (red) LT conditions shown in Fig. 5C are replotted together with the data in 0h LT condition (green) ($n = 54$, pairwise linear correlation, $\rho = -0.5045$, $p < 0.001$).

Table S1

| effect | group | term | estimate | std error | statistic | df | p-value |
|---|---|---|---|---|---|---|---|
| fixed | NA | (Intercept) | -0.03843 | 0.03509 | -1.095 | 5.33 | 0.32 |
| fixed | NA | Lighting condition | -0.03899 | 0.01424 | -2.738 | 17 | 0.014* |
| random | Bird | (Intercept) | 0.07812 | NA | NA | NA | NA |
| random | Bird:Syllable | (Intercept) | 0.04136 | NA | NA | NA | NA |
| random | Residual | Observation | 0.04272 | NA | NA | NA | NA |

Table S2

| effect | group | term | estimate | std error | statistic | df | p-value |
|---|---|---|---|---|---|---|---|
| fixed | NA | (Intercept) | −2.495e-05 | 6.903e-06 | −3.610 | 8.22 | 0.00653 |
| fixed | NA | Lighting condition | 6.737e-06 | 7.009e-06 | 0.96 | 17 | 0.34993 |
| random | Bird | (Intercept) | 1.023e-10 | NA | NA | NA | NA |
| random | Bird:Syllable | (Intercept) | 8.673e-11 | NA | NA | NA | NA |
| random | Residual | Observation | 4.422e-10 | NA | NA | NA | NA |

Table S1 & S2. A linear mixed effects model was run to evaluate whether the LT conditions (−3h LT vs +3h LT) influence the slope of acoustic-distance change in syllable structure during morning singing. Table S1 shows the results of the slopes over the first 30-min period following LT. Table S2 shows the results of the slopes for the first 1000 syllable renditions following LT. In both tables, the outcome variable and the fixed effect were slope and LT condition, respectively. Random effects included random intercepts for bird, and for syllable nested within bird, to account for hierarchical structure and non-independence of renditions.

1. Movie S1. An example of DC-like intensive singing following +3h LT.
2. Movie S2. An example of DC-like intensive singing following -3h LT in the same bird as in Movie S1.
3. Movie S3. Infrared recording of bird movement in the dark before +3h LT, displayed at 16x speed.
4. Movie S4. Infrared recording of bird movement in the dark before +3h LT (the same bird as in Movie S3).
5. Movie S5. Operant conditioning to associate a lever press with lighting.
